# Resource selection at fine-scale: What drives the decision of a generalist herbivore?

**DOI:** 10.1101/2023.12.18.572166

**Authors:** Melinda Boyers, Francesca Parrini

## Abstract

Spatial patterns in topography and forage distribution significantly influence the movements and choices of large herbivores. However, understanding the foraging strategies of free-grazing herbivores at different temporal and spatial scales remains limited, as different behavioural decisions can apply at different hierarchical levels, This study investigates the fine-scale foraging strategies of zebra (*Equus quagga*) in a South African savanna, with a specific focus on their selection of green vegetation at the plant and feeding patch levels. We used the Normalized Difference Vegetation Index (NDVI) as a proxy for vegetation productivity and quality. Our findings reveal that zebra adapted their foraging strategies according to the scale and season. During the late-dry season and early-wet season, selection for greenness was at both the grass tuft and feeding site levels. In contrast, during the mid-dry season, their selection was predominantly at the tuft level, focusing solely on greenness. These insights emphasize the importance of conducting multi-level studies when investigating factors influencing foraging decisions. Findings at one hierarchical level may not necessarily apply across other levels of investigation, highlighting the need for a nuanced and comprehensive approach to understanding the complex foraging behaviours of these animals.

## Introduction

Climate change represents one of the most pressing global challenges, encompassing environmental, social, and economic dimensions Scheffers et al. (2016). Understanding how climate impacts ecosystems remains a challenge within the fields of ecology and natural resource management. A large amount of evidence regarding climate change points to shifts in plant phenology and species distribution (Craine et al., 2009; Piao et al., 2019; Vitasse et al., 2021). Therefore, comprehending the mechanisms such as foraging strategies that free-grazing herbivores adopt at different spatial and temporal scales, becomes a focal point in animal ecology (Balluffi-Fry et al., 2020; Fryxell et al., 2004; Prins, Van Langevelde, 2008) and ecosystem management (Bailey et al., 1996; Senft et al., 1987).

Resource distribution plays a pivotal role in shaping the movement and distribution of herbivores (Muya, Oguge, 2000). Predicting the distribution of large herbivore species, whether for conservation or management purposes, necessitates a clear understanding of how different species utilise available resources. Various factors come into play, including grass height, grass species composition, woody canopy cover ((Ben-Shahar, 1991; Owen-Smith, 2002; Sinclair et al., 1985)), and grass greenness (Mcnaughton, 1985; Sinclair et al., 1985), all of which significantly affect the resources and overall habitat conditions for large grazers. The physical properties and structural characteristics of the grass also impact its acceptability (O’Reagain, Schwartz, 1995). Notably, many animals exhibit a strong preference for green plant material over dry material, as observed in sheep and cattle (O’Reagain, Schwartz, 1995). Furthermore, the greenness of vegetation is inversely correlated with its maturity (Van Soest, 1994), meaning that younger vegetation tends to be greener and in generally indicative of higher-quality forage (O’Reagain, Owen-Smith, 2009).

Spatial patterns in topography and the distribution of forage play a critical role in shaping the movements of large herbivores. However, our understanding of the relative importance of foraging decisions made across different spatial scales remains somewhat limited. Different foraging response patterns are displayed at different scales defining different hierarchies (Bailey et al., 1996; Bowers, 2006; Rettie, Messier, 2000; Senft et al., 1987; Wilmshurst et al., 1999) which may begin at the landscape scale and gradually refines down to finer scales, progressing through feeding patches, feeding stations, and finally, specific plant parts or bites (Bowers, 2006). Decisions made at large temporal and spatial levels, such as where to commence grazing) can significantly influence behaviours at smaller scales, while choices at smaller scales may be integrated and inform decisions at the broader levels (Bailey et al., 1996). Thus herbivores must integrate information from lower levels, including bites, feeding stations, and patches, to assess spatial alternatives at higher levels, encompassing foraging areas, camps, and home ranges (Bailey et al., 1996).

Resource heterogeneity occurs at all spatial scales within the environment, making it challenging to determine in advance at which spatial scale resource selection might predominantly occur (Senft et al., 1987). Different herbivore species exhibit different trade-offs between forage quality (measured by factors like greenness and species) and quantity (forage mass). Zebra, a high-density non-ruminant herbivore, possess a hind-gut digestive system that allows them to process food at a faster rate compared to foregut ruminants, which enables zebra to potentially exploit a wider range of grass quality and quantity (Hack et al., 2002). Previous research in the southwestern region of the Greater Kruger National Park in South Africa (Sabi-Sands Game Reserve and Timbavati Game Reserve) found that grass height and greenness were positively associated with zebra’s preference for specific grasses within their foraging areas (Ben-Shahar, 1991; Bodenstein et al., 2000). In contrast, studies conducted in Punda Maria in Kruger National Park found that zebra accepted a broader range of grass greenness, including brown grass during both the early and late dry seasons (Macandza, 2009). At larger scales, zebra directed their movement towards patches rich in high-quality resources within the expansive natural landscape of Makgadikgadi Pans National Park in Botswana (Brooks, Harris, 2008).

In this study we investigated whether greenness, plant species, or a combination of both factors influenced zebra feeding behaviour across two distinct spatial scales: the grass tuft within feeding stations and feeding stations within foraging areas. The aim is to shed light on how these selection criteria change with scale and across different seasons, contributing to a more comprehensive understanding of zebra feeding behaviour. We expected that within a foraging area, zebra would select feeding stations based on species composition and greenness. Within a feeding station, we expected them to prioritize forage quality by selecting for greenness, while species composition may not be a significant factor in their decision making process at this scale.

## Methods and Materials

### Study Area

This study was conducted in Manyeleti Game Reserve, hereafter referred to as Manyeleti, situated along the western boundary of the Kruger National Park in the Mpumalanga province, close to the border of the Limpopo province in South Africa (Figure 1). Manyeleti is located within the savanna biome, characterised by a discontinuous overstory of woody plants and a herbaceous layer primarily dominated by C_4_ grasses (Venter et al., 2003). The vegetation in this area is an open tree savanna, with the dominant tree species being *Senegalia nigrescens* and *Sclerocarya birrea*. The most dominant shrubs include *Grewia spp, Ziziphus mucronata, Flueggia virosa* and *Ormocarpum trichocarpum*. Within the grass layer, *Heteropogon contortus, Themeda triandra, Panicum maximum* and *Enneapogon spp*. are the dominant species. Various forbs are also present Bodenstein et al. (2000). Manyeleti experiences high average temperatures in summer, with a mean of 32.4 °C, while winters are generally mild, and free from frost, with an average temperature of 17.8 °C. Rainfall in this area is concentrated between October and April, with an annual long-term average of 572 mm over a 41 years long period (Kingfisherspruit, KNP Scientific Services).

**Figure 1.**
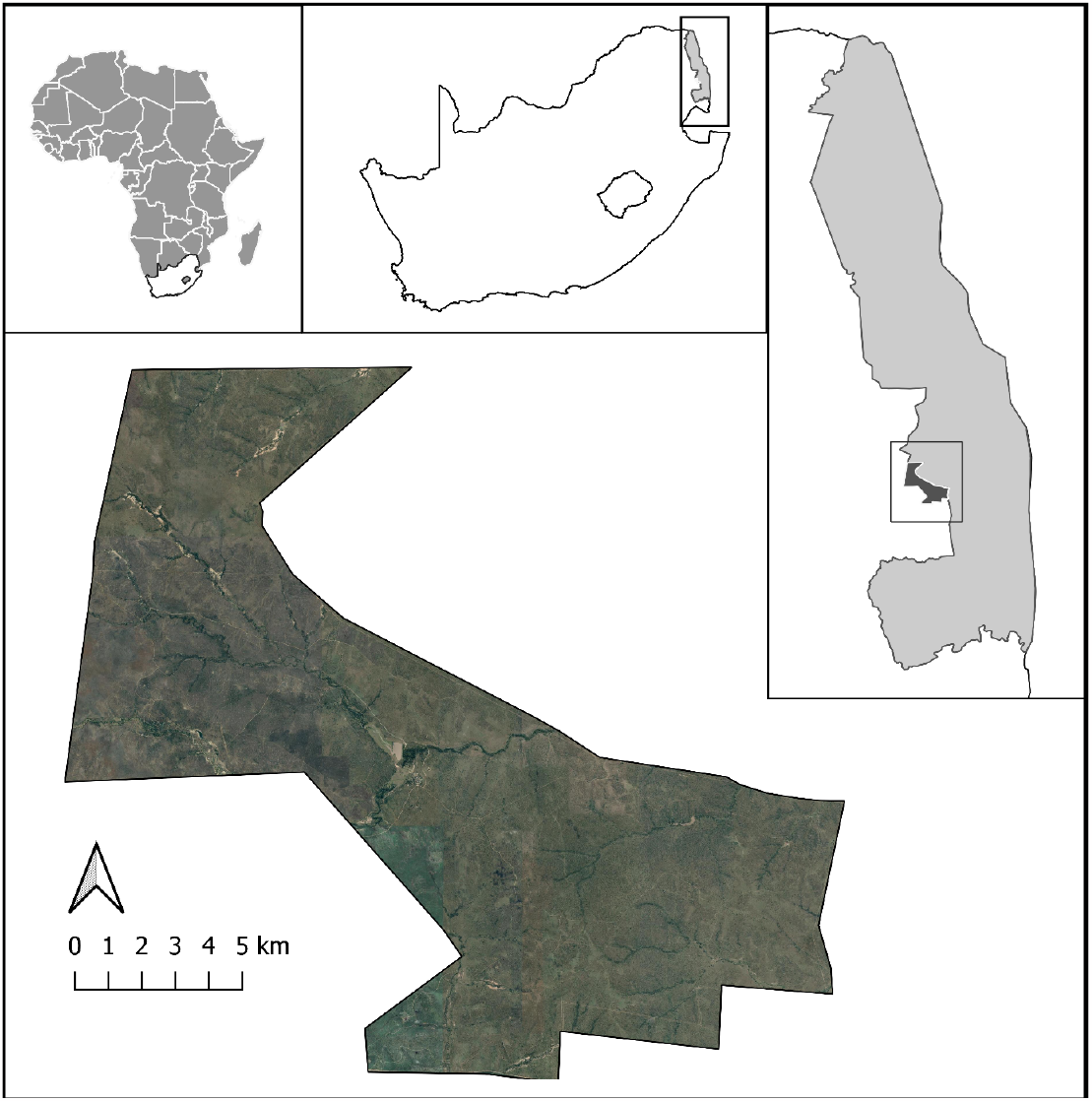
Map of Manyeleti Game Reserve, including continental position (map inset top left), showing the Kruger National Park situated on the border. There are no fences between Manyeleti Game Reserve and the Kruger National Park.

### Field data collection

We sampled data at two spatial scales: the grass tuft within a feeding station and the feeding station within a foraging area. The dry season is a critical period for African ungulates, as the limited rainfall leads to a decline in food quality during this period. To capture this crucial phase of the year, our data collection extended over one dry season, from August to October 2010, and a transition period to the onset of wet season, in November 2010. Within the dry season, we further divided our observations into two distinct periods: the mid-dry season spanning August to September, and the late-dry season in October, based on monthly rainfall and greenness values (Figure 2).

**Figure 2.**
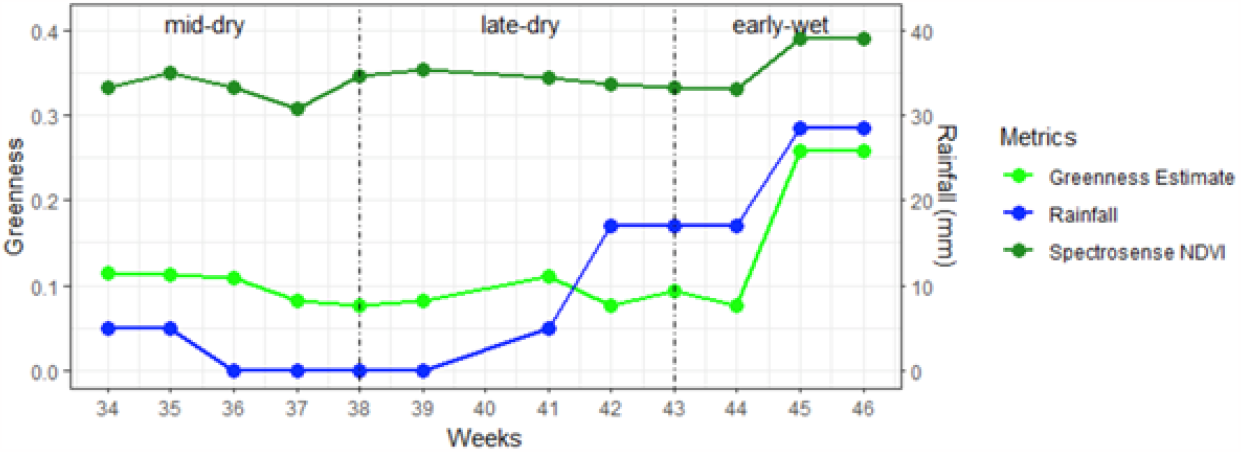
The relationship between the estimated greenness values and SpectroSense NDVI as well as the rainfall over the study period (AUG – NOV 2010).

In the early morning and late afternoon, which are the primary feeding times for ungulates, we drove throughout the reserve to locate grazing zebra individuals or groups within 100 m on either side of the road. To avoid re-sampling the same herd, we did not return to the same area on the same day. Upon identifying a foraging individual or group, we observed their behaviour, and, once they had moved on, approached the foraging area on foot. The location of the foraging area was confirmed by the presence of fresh bites, fresh zebra dung and spoors. Fresh bites are identifiable by the white, undried appearance of the bitten leaf or stem, while older grazed grass turns brown quickly (Kleynhans et al., 2011; Macandza, 2009). Upon identifying the first signs of fresh feeding, we placed a 0.5 m x 0.5 m quadrat to represent a feeding station (Novellie, 1987). We defined a feeding station as the area accessible to the foraging zebra without requiring them to move its legs, consistent with the definitions proposed by Bailey et al. (1996) and Novellie (1987). In cases where fresh footprints and/or signs of cropping revealed a foraging path, we placed nine additional quadrats along the path. If a foraging path could not be identified, we positioned nine additional quadrats surrounding the central one along the four cardinal directions. Grass tufts with fresh bites were classified as “used”, and feeding stations with used grass tufts present were similarly classified as “used”. For every used feeding station we sampled, we also sampled an unused feeding station, which was identified by showing no signs of grazing. These unused feeding stations were randomly positioned, ensuring that they were at least 2 m apart from the used feeding stations. For both used and unused feeding stations we recorded information regarding grass species composition, species basal cover, the season, and the average greenness using a handheld NDVI measuring tool (SpectroSense 2+, Skye Instruments, Netherlands). Within the used feeding stations, we documented both used and unused grass tufts according to species, season, and tuft greenness using Walker (1976) 8-point scale: 0%, 1-10%, 11-25%, 26-50%, 51-75%, 76-90%, 91-99%, 100%.

## Data analysis

We investigated use versus non-use at two distinct spatial levels of selection: feeding stations within a foraging area and grass tufts within a feeding station. We considered a feeding station as “used” when any species present within it displayed clear indications of having been eaten, while a grass tuft was scored “used” if it had fresh bite marks. We categorised our seasons based on rainfall patterns and mean NDVI greenness of the grass. These defined seasons included the mid-dry season (August – September 2010), the late-dry season (October 2010) and the early-wet season (November 2010). For data analysis, we only included the four most abundant grass species that were present in a minimum of 10 foraging areas used by zebra per season. This selection was made to ensure a large enough sample size and enable a reliable comparison of acceptability of these particular grass species. The remainder of the species were collectively categorised as ‘other’.

We analysed our data using Generalized Linear Mixed Effects Models (GLMMs). In our models for feeding station selection, the explanatory variables were presence or absence of individual species, season (mid-dry, late-dry and early-wet), cover, and NDVI values of each feeding station as fixed effects, with foraging area as the random effect. We found collinearity between the basal cover of species and the NDVI measurement. Consequently, species basal cover was excluded from the analysis. In the models addressing grass tuft selection, explanatory variables included grass species, season (mid-dry, late-dry and early-wet), and grass tuft greenness as fixed effects, with foraging area and feeding station as nested random effects. Because of false convergence errors during analysis, when the model fitting functions fail to converge on a maximum likelihood estimate, often due to insufficient data points within a categorical variable level, we combined the greenness ranks 50-75% and 76-90% into a single rank of 50-90% green.

All models were compared using Akaike’s Information Criterion (AIC). Further model comparison was conducted by calculating the relative likelihoods of all the candidate models (wi) and evidence ratios derived from these relative likelihoods (Ei,j = wi/wj). Evidence ratios are used to compare strength of evidence between models within the same set, where a higher evidence ratio signifies stronger support for model i over model j (Anderson, 2008). We also calculated the conditional R^2^ for our mixed-effect models. This metric describes the proportion of variance explained by both the fixed and random factors (Nakagawa, Schielzeth, 2013). For the best model(s), we calculated the log odd ratios from the model coefficients and the 95% confidence intervals from the variances and covariances of the estimated parameters. We used the R statistical software environment (Team, 2018) with R packages lme4 (Bates et al., 2015) to perform the GLMM analysis, AICcmodavg (Mazerolle, 2019) to perform the model selection, and emmeans (Lenth, 2019, 2020) to estimate marginal means from models.

## Results

We sampled a total of 94 foraging areas, 1120 used and non-used feeding stations (in equal numbers) and 4861 grass tufts (2223 grazed and 2638 non-grazed) within the used feeding stations. Throughout or study, we documented a total of 28 different grass species, 9 of which were never eaten by zebra (Appendix I). Of the 19 species consumed, the 4 most abundant species were *Panicum maximum, Urochloa mosambicensis, Themeda triandra, and Digitaria eriantha*. The remaining species were grouped under ‘other’ for data analysis. Our focus in both feeding station selection and grass tuft selection analyses was to discern whether the presence of specific species or their greenness influenced selection. Moreover, we investigated how selection at these spatial scales varied across seasons, specifically the mid-dry, late-dry, and early-wet seasons.

The best model explaining feeding station selection included interactions between feeding station greenness and the presence of species, as well as between seasons and the presence of species (Model 8, AICc weight = 0.72, Appendix II). Specifically, feeding stations with *P. maximum* showed an increased likelihood of being selected as mean greenness increased, up to a greenness to 60%. Beyond this point, the presence of *P. maximum* presence did not further explain selection (Figure 3A). Conversely, feeding stations with *U. mosambicensis* displayed a declining likelihood of selection as mean greenness increased. The remaining species categories (*T. triandra, D. eriantha* and ‘other’) did not increase selection by being present but did increase selection with an increase of mean greenness within the feeding station. In addition, during the late-dry season, there was an increased likelihood of a feeding station being selected with an increase in *P. maximum* and an avoidance of *T. triandra* within the feeding compared to the other seasons (Figure 3B).

**Figure 3.**
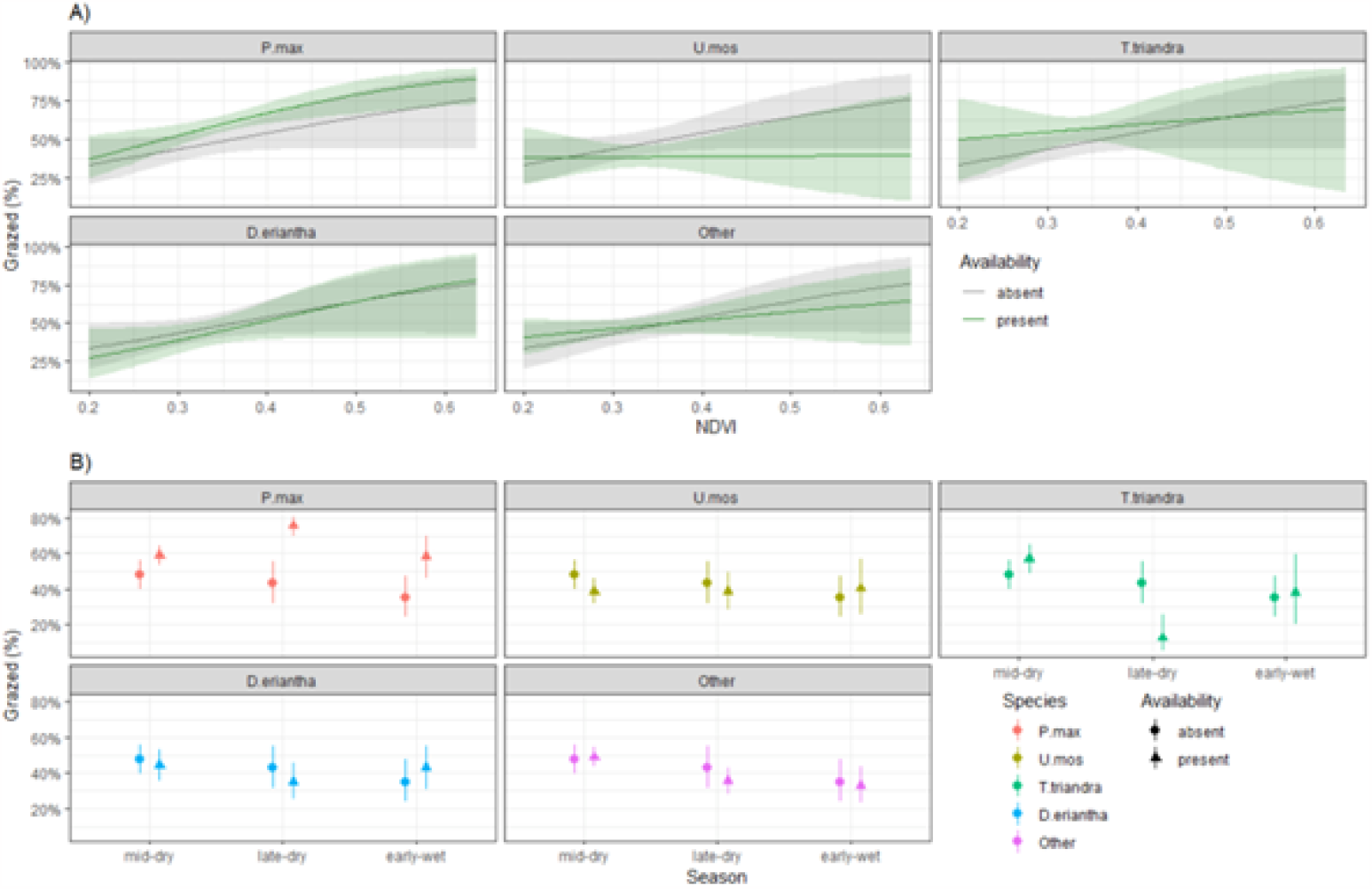
A) Feeding station selection estimates for five species categories within NDVI categories (± 95% confidence interval) for Manyeleti; namely *P*.*max* = *Panicum maximum, U*.*mos* = *Urochloa mosambicensis*, Other = all other grasses, *T*.*triandra* = *Themeda triandra, D*.*eriantha* = *Digitaria eriantha*. the green regression line indicates present and grey regression line represents absent from feeding station. B) Feeding station selection estimates per species per season (± 95% confidence interval) for Manyeleti. From data collected over a three month period between August and November 2010.

The best model explaining within grass tuft selection included the interaction between species and greenness, as well as between species and season (weight and evidence ratio = 1, Appendix III). In both the mid and late-dry seasons, the majority of grass tufts fell below the 10% greenness threshold. The few grass tufts in higher greenness categories were almost 100% grazed when available (Figure 4A). Overall, greenness categories 11-25%, 26-50% and 51-90% were more selected in comparison to those within the 1-10% greenness category (Figure 4B). During the early-wet season, there was an increase in the number of grass tufts within the 11-25% and 26-50% greenness categories, which resulted in zebra grazing more on the greener tufts (Figure 4A). Thus, across all periods, an increase in grass tuft greenness correlated with an increased probability of selection. s. Within each greenness class, when season was held constant, zebra consistently exhibited a preference for *P. maximum* over other species, except in the over 51% greenness category(Figure 4D). Notably, during the late-dry season, *P. maximum* was the preferred grass species while there was no difference in species selection in the other two seasons (Figure 4C).

**Figure 4.**
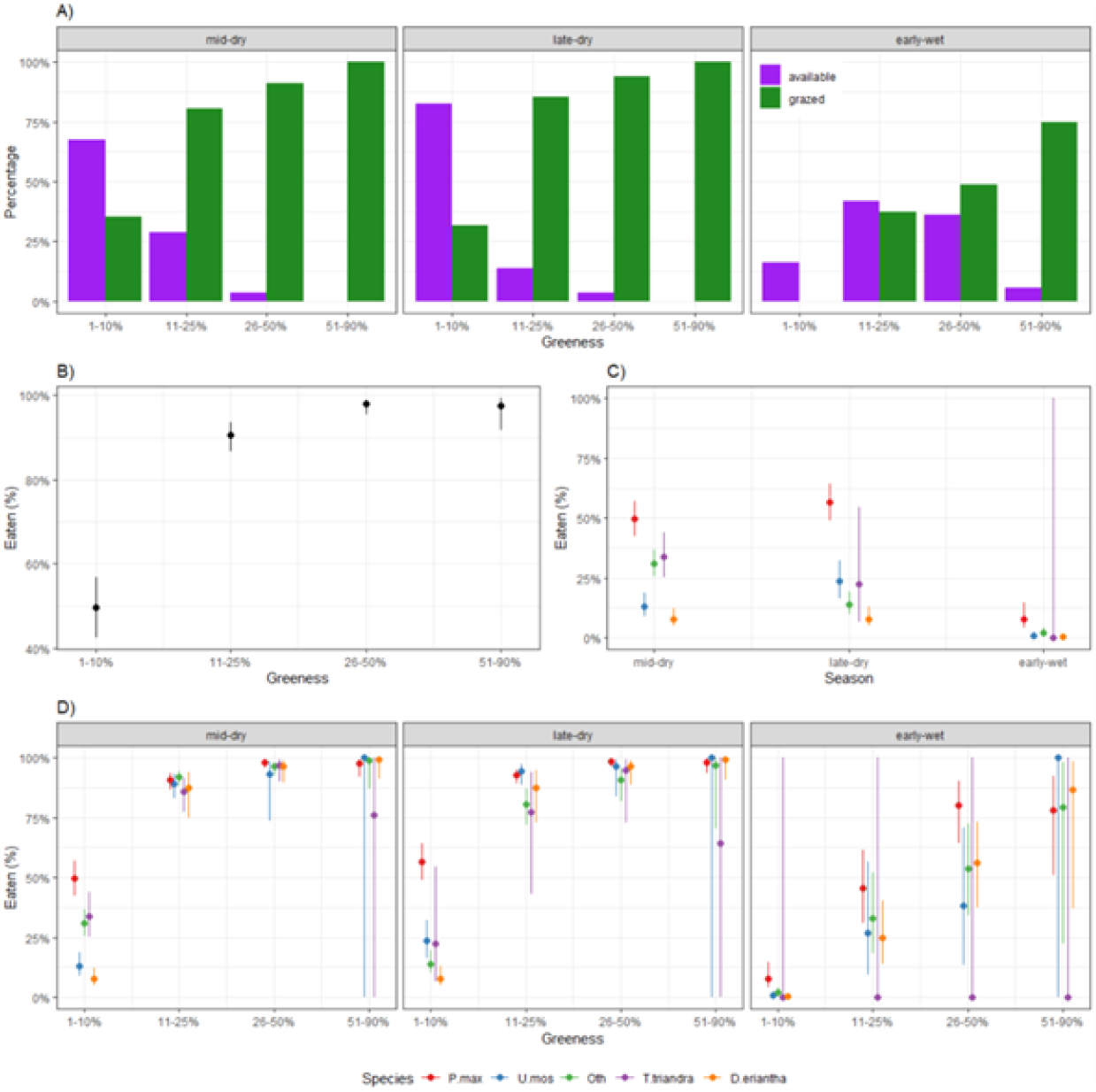
A) Percentage of available grass tufts compared to the percentage of the grass tufts that are grazed within each greenness category in the three different seasons: Mid-dry season, Late-dry season and Early-wet season. B) Grass tuft greenness selection estimates (± 95% confidence interval) for Manyeleti. C) Grass species selection estimates per greenness category (± 95% confidence interval) for Manyeleti. D) Grass species selection estimates per season (± 95% confidence interval) for Manyeleti. *Panicum* = *Panicum maximum, U*.*mos* = *Urochloa mosambicensis*, Other = all other grasses, *T*.*triandra* = *Themeda triandra, D*.*eriantha* = *Digitaria eriantha*. Data collected over a three month period between August and November 2010.

## Discussion

In this study, we analysed zebra selection for feeding stations within foraging areas and grass tufts within these feeding stations. Contrary to our initial expectations, selection for specific species predominantly occurred at the level of individual tufts. Selection for greenness was observed at both levels – feeding stations and grass tufts -during the late-dry season and early-wet season and only at the tuft level during the mid-dry season.

We had hypothesized that zebra would base their selection for feeding stations within foraging areas on both greenness and species composition. Notably, we found that greenness played a significant role in explaining the use of feeding stations only during the late-dry and early-wet seasons, while it did not show a significant effect during the mid-dry season. In contrast, a separate study investigating the foraging area selection of zebra and other grazers in another savanna area of South Africa (Ezemvelo Nature Reserve) did not observe a preference for greenness by zebra during the late-dry season and wet season, but only during the early- and mid-dry season (Mariotti et al., 2020). These discrepancies raise questions about the potential factors contributing to such differences. It is indeed difficult to ascertain if these variations stem from differences in the scale under investigation (within foraging area in our study vs foraging areas within patches; Mariotti et al. (2020)) or if they are attributed to the different environmental conditions present in the two study areas. Our results revealed that although species explained selection for feeding stations in our area, it was mostly in conjunction with greenness. Indeed, feeding stations with *P. maximum* were more likely to be selected, but this preference was observed only when the greenness of the feeding station was above 30%. In instances where greenness was lower, species composition did not explain the selection for feeding stations. Therefore, the selection for feeding stations was not driven by species composition as much as we were expecting, but rather by greenness. The contrasting findings highlight the complexity of factors influencing zebra behaviour and emphasize the importance of considering various factors when interpreting animal foraging decisions in diverse environments.

Within each feeding station, our expectation was that zebra would primarily select grass tufts based on their greenness rather than species. Indeed, while zebra selected grass tufts with the highest greenness, the influence of species was also important. During the late-dry season, characterised by most species displaying low greenness, tufts of *P. maximum* were the most selected among all species. However, an interesting pattern emerged from our results: if zebra had to choose between a tuft of *E. rigidior* with greenness above 51% versus *P. maximum* with greenness less than 10%, they would opt for *E. rigidior*. Interestingly, zebra in the Timbavati Nature Reserve, preferred *U. mosambicensis* over other species including *P. maximum* throughout the year (Bodenstein et al., 2000). In contrast, our study showed a general avoidance of *U. mosambicensis*, except when greenness levels were high. *U. mosambicensis* grows in disturbed and overgrazed areas, which may be the reason for ‘inadvertently’ not selecting feeding stations that presented this species during the mid-dry season. At a broader scale that the ones investigated in this study, Ben-Shahar, Coe (1992) found that zebra exhibited seasonal movement between grass communities characterized by a high proportion of nutritious species, rather than distinct selection for particularly nutritious species within these communities. Our study did not look at the grass community level, but we focused at within foraging area selection, a hierarchical level in between grass tuft and plant community (Senft et al., 1987).

Our findings revealed an interesting trend:, as the feeding stations became greener, the probability that species composition would explain selection decreased. It does indeed make sense that when the overall palatability of the general grass sward increases, selection for specific plant species becomes less pronounced. This suggests that while zebra exhibit selectivity towards certain species, it appears more prominent during the drier months. Studies conducted in other regions, such as Timbavati Nature Reserve (Bodenstein et al., 2000) and Hluhluwe iMfolozi Park (Kleynhans et al., 2011), noted that *P. maximum* made up a significant proportion of the zebra diet of zebra in Timbavati Nature Reserve, similar to our observations in the current study area. *P. maximum* is known for its nutritional value and longer retention of greenness compared to other species, especially during the dry season (Oudtshoorn van, 2009), is preferred by many grazers during that period (Mutanga et al., 2004). Although zebra in our study area favoured *P. maximum* throughout the study duration, during the early-wet season, when most grasses were green, the greenest tufts were selected regardless of species. Our results are in line with previous studies that identified vegetation greenness as the primary driver of herbivore selection (Senft et al., 1987). This consistent pattern emphasizes the significant influence of greenness on herbivore foraging behaviours.

Studies addressing how animals’ selection criteria are influenced by the scale at which the selection operates, have yielded contrasting conclusions. Some studies, such as (Fortin et al., 2005) have demonstrated that different criteria might be used across varying scales, while others, like (Ward, Saltz, 1994), suggest that animals might employ the same selection criteria across different spatial scales. For instance, research on migratory elk (*Cervus elaphus*) revealed distinct forage quality selection patterns at different spatial scales compared to residential elk (Hebblewhite et al., 2008). Migratory elk exhibited different selection patterns at the boarder scales compared to finer scales, while residential elk displayed consistent forage quality selection across all spatial scales. The contrasting selection criteria between migratory and residential elk resulted in improved quality forage for the former but also resulted in heightened predation risk (Hebblewhite et al., 2008). Similarly, our study findings indicate that, like the residential elk, zebra used similar selection criteria across the two spatial scales examined. The primary driving force behind their selection was predominantly attributed to changes in vegetation greenness due to seasonal variations. Although we observed some selection for species composition, the prevalence of *P. maximum*, maintaining its greenness for a longer duration during the dry season, suggests that greenness remains a principal driver of selection. This pattern echoes the findings that greenness plays a pivotal role in influencing animal foraging decisions across different spatial scales.

The relationship between small-scale foraging decisions and the broader distribution patterns of herbivores has been well-documented. At the smaller scales, animals are confronted with spatial variability among the grass canopy, due to inherent variations in the nutrient content among different plant species. Our observations of free-grazing zebra in Manyeleti show that zebra adjust their foraging strategies based on greenness and specific species, particularly at the finer scales. This was notably pronounced during the late-dry season, and somewhat less prevalent in the early-wet season. To deepen our understanding of the foraging strategies of large herbivores, future research should focus on comparing selection dynamics across various hierarchical levels and variations, integrating small scale patterns with broader scales, encompassing foraging areas and beyond, utilizing tools such as satellite remote sensing imagery. This integration could offer a comprehensive perspective, enriching our understanding of how these animals navigate and make foraging decisions across varying scales and changing environmental conditions.

## Acknowledgments

We would like to express gratitude to Mpumalanga Parks and Tourism for accommodating us during this study. A special thanks to Jimmy Thanyani from Manyeleti Game Reserve for access to areas which made this project a success.

## Authors contribution

Both MB and FP contributed to study design, data collection and interpretation. MB wrote the initial draft. All authors approved the final version of the manuscript.

