## Appendices for "Resource selection at fine-scale: What drives the decision of a generalist herbivore?"

generalist herbivore?

Melinda Boyers1*, Francesca Parrini2

1 School of Biodiversity, One Health and Veterinary Medinice, University of

Glasgow, Glasgow G128QQ United Kingdom

2 Centre for African Ecology, School of Animal, Plant and Environmental

Sciences, University of the Witwatersrand, Wits 2050, South Africa

* corresponding:

Supplementary information

Appendix I: List of species sampled within the study area

| Used species | Unused species |
| --- | --- |
| *Panicum maximum*  *Panicum coloratum*  *Urochloa mosambicensis*  *Digitaria eriantha*  *Themeda triandra*  *Heteropogon contortus*  *Setaria sphacelata* var. *Sphacelata*  *Eragrostis superb*  *Eragrostis trichophora*  *Eragrostis rigidior*  *Eragrostis ciliaris*  *Panicum ecklonii*  *Tricholaena monachne*  *Eragrostis lehmanniana*  *Eragrostis chloromelas*  *Cynodon dactylon*  *Chloris virgata*  *Dactyloctenium giganteum*  *Bothriochloa radicans* | *Cymbopogon sp*  *Perotis patens*  *Cenchrus ciliaris*  *Aristida sp*  *Aristida scabrivalvis*  *Sporobulus festivus*  *Sporobulus nitens*  *Pogonarthria squarrosa*  *Sporobulus fimbriatus* |

Appendix II: Some candidate high ranking, and low ranking mixed-effect models, and their coefficients, describing feeding station selection in Manyeleti, showing various explanatory variables and interactions (×) included in models

| Model | Model selection based on AICc: | K AICc | AICc | Delta_AICc | AICcWt | LL |
| --- | --- | --- | --- | --- | --- | --- |
| 8 | NDVI * (T.triandra + U.mos + D.eriantha + P.max + Other) + Season * (T.triandra + U.mos + D.eriantha + P.max + Other) | 25 | 3093.39 | 0 | 0.72 | -1521.4 |
| 5 | Season * (T.triandra + U.mos + D.eriantha + P.max + Other) | 19 | 3095.27 | 1.88 | 0.28 | -1528.4 |
| 6 | NDVI * Season + NDVI * (T.triandra + U.mos + D.eriantha + P.max + Other) | 17 | 3115.88 | 22.48 | 0 | -1540.8 |
| 7 | NDVI * (T.triandra + U.mos + D.eriantha + P.max + Other) | 13 | 3135.91 | 42.51 | 0 | -1554.8 |
| 4 | T.triandra + U.mos + D.eriantha + P.max + Other | 7 | 3136.27 | 42.88 | 0 | -1561.1 |
| 3 | NDVI | 3 | 3243.99 | 150.6 | 0 | -1618.9 |
| 1 | only intercept | 2 | 3259.01 | 165.61 | 0 | -1627.5 |
| 2 | Season | 4 | 3263.01 | 169.61 | 0 | -1627.5 |

Appendix III: Candidate mixed-effect models, and their coefficients, describing grass tuft selection in Manyeleti, showing various explanatory variables and interactions (×) included in models listed from higher to lower ranked.

| Model selection based on AICc: | K | AICc | Delta_AICc | AICcWt | Cum.Wt | LL |
| --- | --- | --- | --- | --- | --- | --- |
| Species * Greenness + Species * Season | 37 | 5159.51 | 0 | 1 | 1 | -2542.46 |
| Greenness * Species + Season | 29 | 5198.55 | 39.04 | 0 | 1 | -2570.1 |
| Species * Greenness + Season * Greenness | 35 | 5202.13 | 42.63 | 0 | 1 | -2565.81 |
| Species + Greenness + Season | 13 | 5230.6 | 71.1 | 0 | 1 | -2602.26 |
| Species * Greenness | 27 | 5264.31 | 104.81 | 0 | 1 | -2605 |
| Species + Greenness | 11 | 5292.86 | 133.36 | 0 | 1 | -2635.4 |
| Greenness + Season | 9 | 5510.75 | 351.25 | 0 | 1 | -2746.36 |
| Greenness * Season | 15 | 5515.09 | 355.58 | 0 | 1 | -2742.5 |
| Greenness | 7 | 5577.97 | 418.46 | 0 | 1 | -2781.97 |
| Species * Season | 17 | 6177.71 | 1018.21 | 0 | 1 | -3071.79 |
| Species + Season | 9 | 6236.07 | 1076.56 | 0 | 1 | -3109.02 |
| Species | 7 | 6245.56 | 1086.05 | 0 | 1 | -3115.77 |
| Season | 5 | 6609.76 | 1450.25 | 0 | 1 | -3299.87 |
